# High nutritional conditions influence feeding plasticity in *Pristionchus pacificus* and render worms non-predatory

**DOI:** 10.1101/2024.08.27.609904

**Authors:** Veysi Piskobulu, Marina Athanasouli, Hanh Witte, Christian Feldhaus, Adrian Streit, Ralf J. Sommer

## Abstract

Developmental plasticity, the ability of a genotype to produce different phenotypes in response to environmental conditions, has been subject to intense studies in the last four decades. The self-fertilizing nematode *Pristionchus pacificus* has been developed as a genetic model system for studying developmental plasticity due to its mouth-form polyphenism that results in alternative feeding strategies with a facultative predatory and a non-predatory mouth form. Many studies linked molecular aspects of the regulation of mouth-form polyphenism with investigations of its evolutionary and ecological significance. Also, several environmental factors influencing *P. pacificus* feeding structure expression were identified including temperature, culture condition and population density. However, the nutritional plasticity of the mouth form has never been properly investigated although polyphenisms are known to be influenced by changes in nutritional conditions. For instance, studies in eusocial insects and scarab beetles have provided significant mechanistic insights into the nutritional regulation of polyphenisms but also other forms of plasticity. Here, we study the influence of nutrition on mouth-form polyphenism in *P. pacificus* through experiments with monosaccharide and fatty acid supplementation. We show that in particular glucose supplementation renders worms non-predatory. Subsequent transcriptomic and mutant analyses indicate that *de novo* fatty acid synthesis and peroxisomal beta-oxidation pathways play an important role in the mediation of this plastic response. Finally, the analysis of fitness consequences through fecundity counts suggests that non-predatory animals have an advantage over predatory animals grown in the glucose-supplemented condition.

**Research highlights:** This study represents the first systematic attempt to investigate the influence of nutrition on mouth-form polyphenism in the genetic model organism *Pristionchus pacificus*. Through glucose and oleic acid supplementation we show that high nutritional conditions influence feeding plasticity and render worms non-predatory. Mutant analysis indicates a role of *de novo* fatty acid synthesis and peroxisomal beta-oxidation pathways for these responses.

## Introduction

Developmental plasticity allows organisms to alter their developmental trajectory to execute distinct phenotypes in response to environmental cues with polyphenisms representing the most remarkable form of plasticity by exhibiting environment-sensitive alternative phenotypes without intermediate forms (West-Eberhard, 2003). In general, developmentally plastic responses can be observed at physiological, morphological, and behavioural levels, including life history traits of organisms. Many case studies have shown developmental plasticity to be ubiquitous in nature. It has been suggested to facilitate adaptation to novel or heterogenous environments and to play an important role for evolutionary diversification and the creation of novelty (Ghalambor et al., 2007; Pigliucci, 2001; West-Eberhard, 2003). Multiple environmental factors have been shown to influence polyphenisms including temperature, seasonal changes, and population density (see reviews; Simpson et al., 2011; Yang & Pospisilik, 2019). Specifically, nutrition is an important regulator of polyphenisms. Several studies in insects have highlighted the importance of nutritional conditions during development for the generation of discrete phenotypes, revealing genetic and epigenetic mechanisms involved (Casasa et al., 2020; Chandra et al., 2018; Gotoh et al., 2014; Kamakura, 2011; Kucharski et al., 2008). For instance, caste polyphenism in honeybee and horn polyphenism in dung beetles represent some of the extreme cases of nutrition-induced developmental plasticity (Emlen, 1997; Haydak, 1970; Moczek, 1998). Nonetheless, our understanding of how nutrition affects the expression of polyphenic traits in other taxa remains limited.

Besides insects, nematodes provided substantial mechanistic insights into the regulation of polyphenisms in the last decade. In particular, the free-living hermaphrodite *Pristionchus pacificus* has been a model system for the studies of developmental plasticity. Research in *P. pacificus* has provided insights into the genetic and epigenetic regulation as well as the ecological and evolutionary significance of plasticity (Schroeder, 2021; Sommer, 2020). In nature, *P. pacificus* can be found in soil and on scarab beetles (chafers, stag beetles, dung beetles), which, after their death, provide a nutritious environment for the nematode through proliferation of bacteria and fungi on the carcass (Herrmann et al., 2006; Meyer et al., 2017; Ragsdale et al., 2015; Renahan et al., 2021; Renahan & Sommer, 2022). In contrast to *Caenorhabditis elegans* and most other free-living nematodes, *P. pacificus* shows a remarkable mouth-form dimorphism, which influences its dietary niche and behaviour. During postembryonic development, genetically identical *P. pacificus* worms form one of two, alternative morphs, the so-called eurystomatous (Eu) and stenostomatous (St) mouth forms (Figure 1a) (Bento et al., 2010). While the Eu morph enables facultative predatory feeding on other nematodes by exhibiting two “teeth” and a wide mouth, the St morph leads to strict microbial feeding and forms only one tooth and a narrow mouth (Theska et al., 2020). Major components of the gene regulatory network that govern mouth-form plasticity have been identified in this model (Bui et al., 2018; Casasa et al., 2023; Levis & Ragsdale, 2023; Namdeo et al., 2018; Ragsdale et al., 2013; Serobyan et al., 2016; Sieriebriennikov et al., 2018, 2020; Werner et al., 2023). For instance, a supergene locus with the sulfatase *eud-1* and two alpha-N-acetylglucosaminidase genes (*nag-1* and *nag-2*), as well as the sulfotransferase, *seud-1*/*sult-1*, act as part of a main switch in mouth-form determination (Bui et al., 2018; Namdeo et al., 2018; Ragsdale et al., 2013; Sieriebriennikov et al., 2018). They are involved in the regulation of the evolutionarily conserved nuclear hormone receptors *nhr-40* and *nhr-1* that have been co-opted for the regulation of mouth-form plasticity and act downstream of *eud-1* and *seud-1/sult-1* (Sieriebriennikov et al., 2020; Theska & Sommer, 2024). It is important to note that the understanding of the molecular regulation of feeding structure polyphenism is far from completion with many epigenetic aspects still to be identified (Brown et al., 2024; Werner et al., 2023).

**Figure 1.**
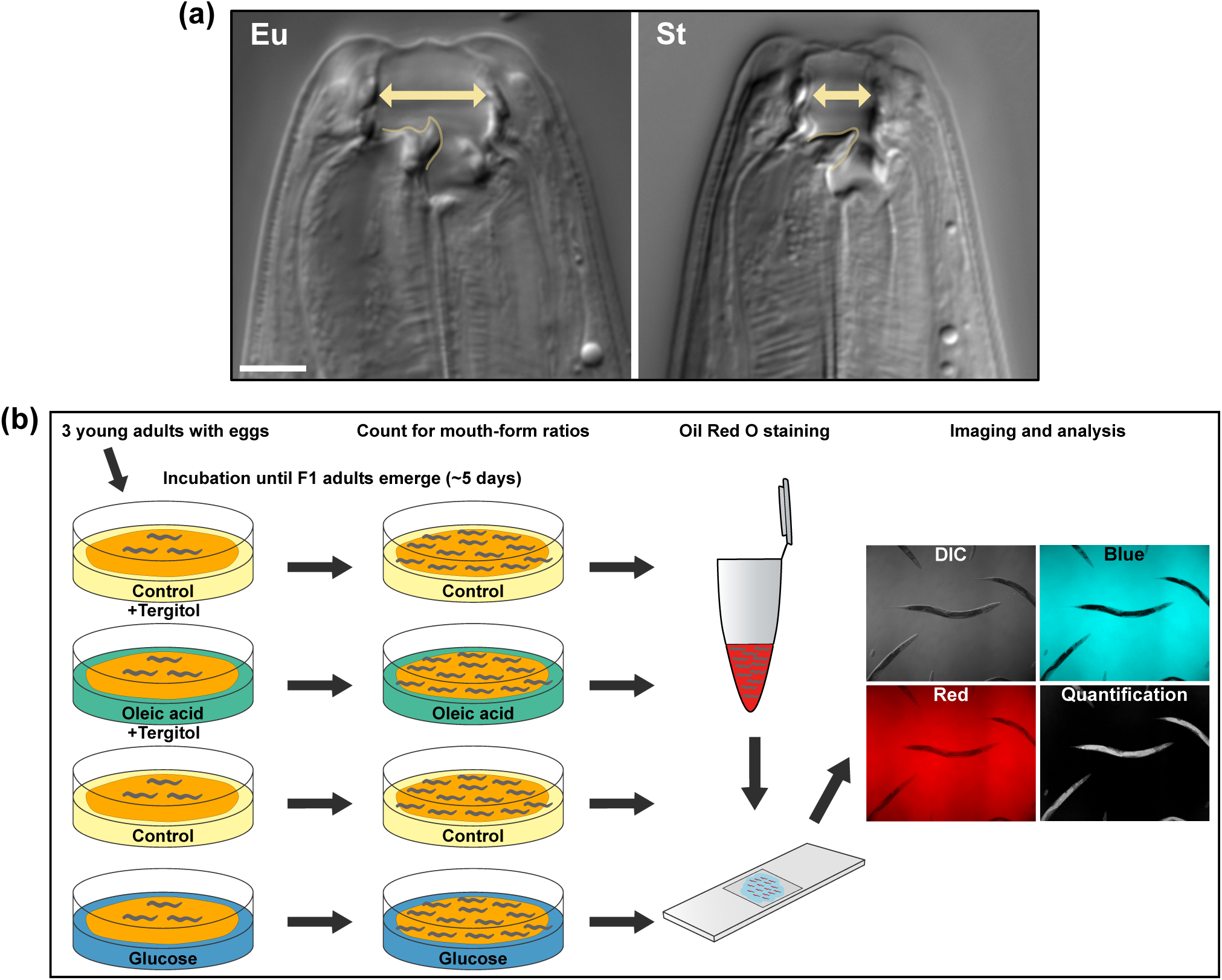
Mouth-form polyphenism of *P. pacificus*, and the experimental design for supplementation studies. (a) Differential interference contrast (DIC) images of the eurystomatous (Eu) and the stenostomatous (St) mouth morphs. The Eu morph has a wide buccal cavity, a hook-like dorsal tooth (on the left, outlined), and a sub-ventral tooth. The narrow-mouthed stenostomatous (St) morph has a flint-shaped dorsal tooth (on the left, outlined). Scale bar is 5μm. (b) Illustration of the general experimental design using supplementation with monosaccharides and fatty acids.

Mouth-form plasticity in *P. pacificus* is influenced by several environmental factors such as culture condition, population density, and temperature (Lenuzzi et al., 2023; Werner et al., 2017, 2018). Previous studies have also revealed prominent roles for nematode metabolites (e.g. ascarosides) and dietary bacteria in the mouth-form decision and predatory feeding (Akduman et al., 2020; Bose et al., 2012; Dardiry et al., 2023). In contrast, our knowledge of how changes to the nutritional status of worms affect mouth-form plasticity is still scarce. Bento et al. (2010) have shown that starvation promotes the development of the Eu morph. More recent genetic work by Casasa and co-workers identified the Mediator subunit MDT15/MED15 to regulate mouth-form polyphenism, which is known to have a conserved role in metabolic responses to nutritional stress (Casasa et al., 2023). These observations suggests that mouth-form plasticity is sensitive to nutritional conditions in particular to low nutrition. However, mouth-form and fitness consequences of a high nutritional condition have never been investigated in *P. pacificus*. One approach of studying the effect of nutrition on nematode plasticity is to introduce supplements, such as monosaccharides and fatty acids, into the standard diet to increase fat storage in nematodes, which is a strong alteration to the nutritional status. This aided research in *C. elegans* to discover novel functions of molecular factors involved in lipid metabolism and associated biological processes (Alcántar-Fernández et al., 2018; Han et al., 2017; Lee et al., 2015; Nomura et al., 2010; Wan et al., 2022). Note that while nutrient supplementation is well established in *C. elegans*, the role of direct and indirect effects of supplementation through the bacterial diet cannot be disentangled. Very likely, the effects observed in such experiments are due to both direct and indirect uptake of nutrients. Moreover, nutrient supplementation studies have prioritised investigating lipid storage as an indicator of nutritional status. Nematodes lack an adipose tissue for lipid storage; however, they generate lipid droplets in many cells including the intestine to serve this function. In addition, storage lipids can be visualised through staining methods such as Oil Red O (ORO), which is a reliable post-fix technique, providing a proxy for lipid storage when it is quantitatively measured (O’Rourke et al., 2009).

Here, we study the influence of nutrition on mouth-form plasticity in *P. pacificus* by establishing experimental setups in which worms are grown in “high nutrition” dietary conditions that promote fat storage. These conditions are obtained through supplementation of monosaccharides and fatty acids (mainly glucose and oleic acid) into the nematode growth medium (NGM) (Figure 1b). We also examine the lipid storage in worms via ORO staining and imaging analysis. We show that fat storage-promoting conditions, in particular glucose supplementation, render worms non-predatory. Similarly, we test whether these dietary effects are mediated transgenerationally but find no such influence. In addition, we carry out transcriptomic and subsequent mutant analyses to explore associated molecular mechanisms. Results indicate that *de novo* fatty acid synthesis and peroxisomal beta-oxidation pathways, which are involved in storage and utilisation of lipids, are essential for nutrition-induced mouth-form plasticity. Finally, we examine worm fitness by using fecundity as proxy to explore whether the mouth-form response in the glucose-supplemented dietary condition benefit *P. pacificus*. Our findings suggest that non-predatory animals have a fitness advantage over predatory animals grown under the same nutritional condition. Overall, this work establishes nutritional status as an additional factor influencing mouth-form polyphenism and highlights the beneficial advantage of this plastic ability in *P. pacificus*.

## Materials and methods

### Nematode stock maintenance

All *P. pacificus* strains were cultured on 6cm NGM plates with 300μl *E. coli* OP50 provided as food at 20°C as previously described (Sommer et al., 1996). The wild type strain PS312 was used for experimentation unless otherwise noted. Stocks were maintained by passing three young adult hermaphrodites with eggs every five days. Mutant *P. pacificus* strains were similarly cultured and are available from the SommerLab.

### Monosaccharide supplementation experiments

Monosaccharides were purchased from Sigma-Aldrich as follows: glucose (D-(+)-Dextrose, product number: D9434), galactose (D-(+)-Galactose, product number: G0750), fructose (D- (−)-Fructose, product number: F0127). All monosaccharides were dissolved in purified and autoclaved water to obtain 1M stock concentrations. Stock solutions were filtered through 0.22μm sterile filters before storage. Plate pouring was carried out under a sterile hood. Each monosaccharide stock was diluted into liquid NGM media to obtain desired final concentrations. Monosaccharide-added NGM was then stirred for approximately 1 minute to obtain a homogenous media. Media was then poured into 6cm plates using a sterile 10ml serological pipette, obtaining a volume of 9ml in each plate. Control plates with the same volume were poured from the main NGM media without monosaccharides. Plates were allowed to dry for 2 days and were then seeded with 300μl overnight grown *E. coli* OP50. After two days, three young hermaphrodites with eggs (from stocks grown under standard conditions) were added onto each plate for each strain, condition, and replicate. After adding worms, plates were incubated at 20°C until F1 adults emerged. Plates were then used for mouth-form phenotyping and ORO staining.

### Fatty acid supplementation experiments

Fatty acid supplementation was carried out as described by Deline et al. (2013); however, cholesterol was added into the NGM media after autoclaving. Sodium oleate (Sigma-Aldrich, product number: O7501) and sodium linoleate (Sigma-Aldrich, product number: L8134) were used for oleic acid and linoleic acid supplementation, respectively. For each experiment, a 0.1M fatty acid working stock was prepared fresh in purified and autoclaved water. Once completely dissolved, the fatty acid working stocks were purged with nitrogen gas to prevent oxidation by air. Plate pouring was carried out as in monosaccharide supplementation except the volume of the NGM in each plate was 8ml. Both supplemented and control plates contained 0.1% Tergitol (NP-40, Sigma-Aldrich, product number: NP40S) to facilitate fatty acid absorption. Once poured, plates were covered with a box to avoid light oxidation. The same steps as in monosaccharide supplementation were followed for seeding the plates with *E. coli* OP50, culturing nematodes, mouth-form phenotyping and ORO staining.

### Mouth-form phenotyping

Mouth-form phenotypes were scored using Zeiss Discovery V.20 stereomicroscope based on mouth-width of individual worms as previously described (Ragsdale et al., 2013; Theska et al., 2020). Each plate was taken as a biological replicate and 30 worms were scored for their mouth-form, unless otherwise mentioned. The percentage of Eu animals of each plate was then calculated. Mouth-form percentage (Eu%) is graphed as a mean of percentages of all plates for each strain in a particular dietary condition.

### Transgenerational experiments

For testing potential transgenerational effects, lines were established by picking three young (day 1) adult worms with eggs from standard condition onto oleic acid, glucose, and control plates, respectively. Ten lines were established for each condition and each line was propagated by transferring three young adult worms for five generations on the same condition. In case of contamination, the line was discontinued and not used for counting mouth form or reversal. At every generation, 6-7 lines (among the already established 10 lines) were selected and reverted back to standard culture conditions. For each condition, mouth-form phenotypes were scored for all plates at every generation and reversal.

### Oil Red O staining

Post-fix ORO staining method was modified from Li et al. (2016). Oil Red O stock solution was prepared by dissolving 0.5g ORO (Sigma-Aldrich, product number: O0625) in 100ml isopropanol. This stock solution was shaken on a see-saw rocker for several days. To prepare ORO working solution, required amount was taken from the stock and centrifuged for 5 minutes at 4500rpm to pellet all ORO-related precipitates. Working ORO solution was prepared fresh by diluting precipitate-free ORO (supernatant) in purified water to obtain 60% ORO. This solution was also shaken on a see-saw rocker for 10 minutes and centrifuged for 5 minutes at 4500rpm before staining worms. For worm fixation, a 1% formaldehyde solution was used by diluting 1ml 16% formaldehyde solution (Thermo Scientific, catalogue number: 28906) in 15ml 1X PBS (phosphate buffered saline). For dehydrating worms, 60% isopropanol was prepared fresh by diluting isopropanol in purified water. First, worms were washed off the plates with 1X PBS and collected in a 15ml Falcon tube for each sample. Note that, after mouth-form phenotyping, several biological replicates (plates) were washed into one sample tube for each strain for a particular dietary condition. Once worms formed a sediment, extra volume was removed, leaving only 1ml. This volume containing worms was then transferred to a new 1.5ml tube. To pellet worms after transfer or wash, samples were briefly centrifuged in a quick-spin centrifuge. Worms were fixed with 1% formaldehyde (1ml) for 30 minutes by shaking on a see-saw rocker. Then, samples were frozen in liquid nitrogen (∼8 seconds) and thawed under running tap water, three times in total. Samples were washed with 1X PBS three times and were dehydrated in 60% isopropanol (1ml) for 2 minutes. Finally, samples were stained with 0.5ml 60% ORO working solution for 30 minutes on a see-saw rocker. The staining solution was applied through a 0.22μm sterile filter. After staining, samples were washed three times with 1X PBS. Supernatant of each sample was removed, leaving approximately 0.1ml sample of stained worms. Two drops of Vectashield (Biozol, catalogue number: H-1000) was then added into each sample. Samples were gently mixed by pipetting up and down before mounting onto microscope slides. Samples were then imaged for ORO quantification.

### ORO quantification and analysis

Oil Red O quantification was performed as described in Feldhaus and Piskobulu (2024 preprint). Briefly, we imaged worms on an AxioImager.Z1 (Zeiss, Oberkochen, Germany) with a 10x/0.3 objective and an AxioCam 506 mono. For ORO quantification, images were obtained with filter sets for DAPI and Texas Red in transmission mode. The filter-based imaging with a monochrome camera was chosen to remove any potential bias from white balance settings of the camera. Oil Red O absorbance per worm was semi-automatically calculated by manually outlining the worm area and then obtaining the difference between the Texas Red channel (where ORO appears transparent) and the DAPI channel (where ORO absorbs) after correcting for differences in the spectral response for both channels. In total, 15 adult worms were quantified per strain and condition. For each worm, raw integrated density (RawIntDen), from channel 4 of the processed image, was taken as a measure of ORO absorbance. To compare individuals with different body size (e.g. mutants), ORO absorbance (RawIntDen) was normalised by the body area (μm^2^) for each worm (RawIntDen/body area). For calculating body area, images of worms from the blue channel (DAPI) were saved in grayscale. The body area was then calculated for each worm by “Threshold” and “Analyze Particles” functions on Fiji (Schindelin et al., 2012). Therefore, the final ORO absorbance value was determined as RawIntDen divided by body area for each worm.

### RNA extraction

For RNA extraction, three young adults with eggs, from standard condition, were placed on oleic acid, glucose, and control conditions, respectively. For each condition, we scanned through 5-9 plates and collected 150 F1 young adults without eggs in two separate tubes for RNA extraction (two technical replicates). RNA extraction was carried out using Direct-zol RNA Microprep Kit (Zymo Research, R2060). Quality of RNA was assessed using NanoDrop. Concentration of RNA was assessed using Qubit 2.0 Fluorometer (Invitrogen). Samples were then diluted to required concentration and sent to NovoGene Co., Ltd for sequencing.

### RNA-seq and KEGG enrichment analysis

Raw reads were aligned to the reference *P. pacificus* genome (El Paco) with STAR (version 2.7.1a) and quantified with featureCounts from the Subread R package (version 2.0.1) based on the latest gene annotations (Athanasouli et al., 2020; Dobin et al., 2013; Liao et al., 2014). The count matrix was filtered by removing genes with less than 10 reads, reducing the number of genes to be examined from 28896 to 18174 and 18049 for oleic acid and glucose respectively. Differential gene expression analysis was performed for each of the two conditions with DESeq2 (FDR-corrected P<0.05, version 1.42.0) (Love et al., 2014). The differentially expressed genes from each condition were tested for overrepresentation of KEGG pathways using Fisher’s exact-test with multiple testing correction (Bonferroni corrected P<0.05) in R (version 4.3.1), based on the existing KEGG annotations for the *P. pacificus* genes (Athanasouli et al., 2023).

### Phylogenetic tree

All *C. elegans* fatty acid desaturase protein sequences were obtained from WormBase (www.wormbase.org). Longer isoforms were picked for genes with different isoforms. Each *C. elegans* fatty acid desaturase protein was then blasted against *P. pacificus* on the protein database (www.pristionchus.org, El Paco annotation v3, 2020). From each blast result, all *P. pacificus* hits were collected and analysed on InterProScan (www.ebi.ac.uk) for conserved protein domains. We found two genes (PPA03289 and PPA05783) which did not contain desaturase domains, which were discarded. The rest of the protein sequences were aligned by ClustalW on MEGA11 software (Tamura et al., 2021). Based on this alignment, a maximum likelihood phylogenetic tree was constructed with LG model, and 100 bootstrap replications. The phylogenetic tree was visualised and edited on FigTree software (version 1.4.4).

### CRISPR/Cas9 mutagenesis and identification of mutants

For the generation of mutants by CRISPR, the protocols of Witte et al. (2015) and Han et al. (2020) were followed with minor modifications. All components of the CRISPR/Cas9 complex were purchased from Integrated DNA Technologies (IDT). Each CRISPR RNA (crRNA) was designed to target 20bp upstream of the protospacer adjacent motif (PAM). To construct CRISPR/Cas9 complex, 5μl CRISPR RNA (crRNA) from 100μM stock was mixed with 5μl trans-activating CRISPR RNA (tracrRNA) obtained from 100μM stock (catalogue number 1072534). This mixture was first denatured at 95°C for 5 minutes, and then cooled down at room temperature for annealing (5 minutes). Hybridized crRNA:tracrRNA product (5μl) was mixed with 1μl Cas9 protein from 62μM stock (catalogue number: 1081058) and incubated at room temperature for 5 minutes. Tris-EDTA buffer was then added into the CRISPR/Cas9 mix to obtain a final concentration of 18.1μM for crRNA:tracrRNA hybrid and 2.5μM for Cas9. Co-injection marker, with *Ppa-eft-3* promoter and TurboRFP sequences, was also integrated into the injection mix (Han et al., 2020). Injected worms were allowed to lay eggs for 24 hours. Fluorescent marker-positive plates were used to isolate emerging F1 progeny. From each positive plate, 8 -10 F1 worms were isolated. Each F1 worm was allowed to lay eggs and then was lysed in single worm lysis buffer (10mM Tris–HCl at pH 8.3, 50mM KCl, 2.5mM MgCl2, 0.45% NP-40, 0.45% Tween 20, 120μg/ml Proteinase K) in a thermal cycler program with 65°C for 1 hour; 95°C for 10 minutes. Lysates were used to carry out polymerase chain reactions to amplify the target region. Mutants in F1 generation (predominantly heterozygotes) were first identified through Sanger sequencing (GENEWIZ Germany GmbH). Subsequent re-isolation and sequencing of the following generation (F2) allowed capturing homozygous mutants. All crRNAs, and related primers for target genes are provided in Table S1.

### Genetic repair of *Ppa-pddl-1(tu2028)*

To genetically revert *Ppa-pddl-1(tu2028)* back to wild type through CRISPR editing, a crRNA (5’-TACTATCCTCTATCCTATAG-3’) was designed targeting 20bp upstream of the TGG PAM site at exon 4 based on the mutant sequence including 7bp insertion. A 100bp wild type repair template, covering the target region, was also used (synthesised by IDT). The repair template was as follows: 5’-CATGAATAGTTCTAACCACTTCTTCTCCTTCAGACATTACTATCCTCTAGTGGCCCTTCTCTGTTTCCTCATGCCAACTGTCGTTCCTGTCTATTATTGG-3’. The repair template was integrated into the CRISPR injection mix at a concentration of 10μM. Homozygous wildtype worms were isolated in F2 generation post injection, validated via sanger sequencing.

### Fecundity experiment

To study fecundity of Eu and St morphs, young adult worms without eggs (F1) were isolated from glucose-supplemented diets (80mM concentration) based on their mouth morph and subsequently transferred to standard dietary conditions with 20μl *E. coli* OP50. Worms were passed onto new plates for 8 consecutive days and were removed at the end of 8^th^ day. After five days from inoculation, viable progeny was counted on each plate.

### Data visualisation and statistical analysis

All plots were generated in RStudio (version 1.2.5042) using ggplot2 and ggpubr packages. Illustrations were produced in Affinity Designer (version 1.9.2). Statistical significance testing was carried out by comparing two independent groups, e.g., control vs. treatment, mutant vs. wildtype, Eu vs. St, for ORO absorbance and total fecundity. Briefly, data was first tested for normal distribution mostly via Shapiro-Wilk normality test with additional visual observations. In case of normal distribution, either two sample t-test (equal variance) or Welch two sample t-test (unequal variance) were used. In case of nonnormal distribution, Wilcoxon rank sum-test was used as the nonparametric equivalent. All statistics were performed in RStudio (version 1.2.5042).

## Results

### Monosaccharide and fatty acid supplementations render worms non-predatory in a concentration dependent manner

For all experiments, we studied hermaphrodites of the PS312 strain, the wild type of *P. pacificus* that had its genome being sequenced at chromosome level (Dieterich et al., 2008; Rödelsperger et al., 2017). This strain is highly Eu under standard lab conditions on an *E. coli* OP50 diet with more than 95% of animals expressing the Eu morph. First, we performed pilot experiments with monosaccharides ranging from 20mM to 300mM concentrations. Results revealed a concentration dependent effect on mouth-form plasticity. Specifically, the number of St worms increased with increasing concentrations of monosaccharides (Figure S1a). Among all, we prioritised glucose as the main St-inducing monosaccharide and performed further experiment by using 100mM and 150mM concentrations. Results indicated that 100mM glucose is sufficient to render worms highly St without exerting observable adverse effects (Figure 2a). Note that even under glucose-supplementation conditions, there is still an effect of population density on mouth-form expression as pilot experiments starting from 10 adult worms (Figure S1a) revealed higher Eu ratios than similar glucose concentration experiments with three adult inoculations (Figure 2a). For fatty acid supplementations, we used oleic acid (monounsaturated fatty acid) and linoleic acid (polyunsaturated fatty acid) to test 0.5mM and 1mM concentrations. Similar to monosaccharide supplementations, oleic acid effect on mouth-form plasticity was mediated in a concentration dependent manner (Figure 2a). Observations based on morphology and developmental pace of worms at both concentrations suggested that 0.5mM oleic acid would be more ideal for further experiments. Results also indicated that the same concentrations were detrimental for worms in the linoleic acid condition. While worms did not populate the plates at 1mM, they were developmentally slow and almost completely St at 0.5mM linoleic acid (Figure S1b). Therefore, we performed another experiment for linoleic acid with reduced concentrations ranging from 0.01mM to 0.2mM. We found that 0.2mM linoleic acid renders worms predominantly St, alleviating the detrimental effect observed at 0.5mM concentration (Figure S1c). Overall, these results show that monosaccharides and fatty acid supplementations induce changes of the mouth-form ratio towards St in *P. pacificus*. For further experiments, we selected oleic acid and glucose as main supplements at 0.5mM and 100mM concentrations, respectively.

**Figure 2.**
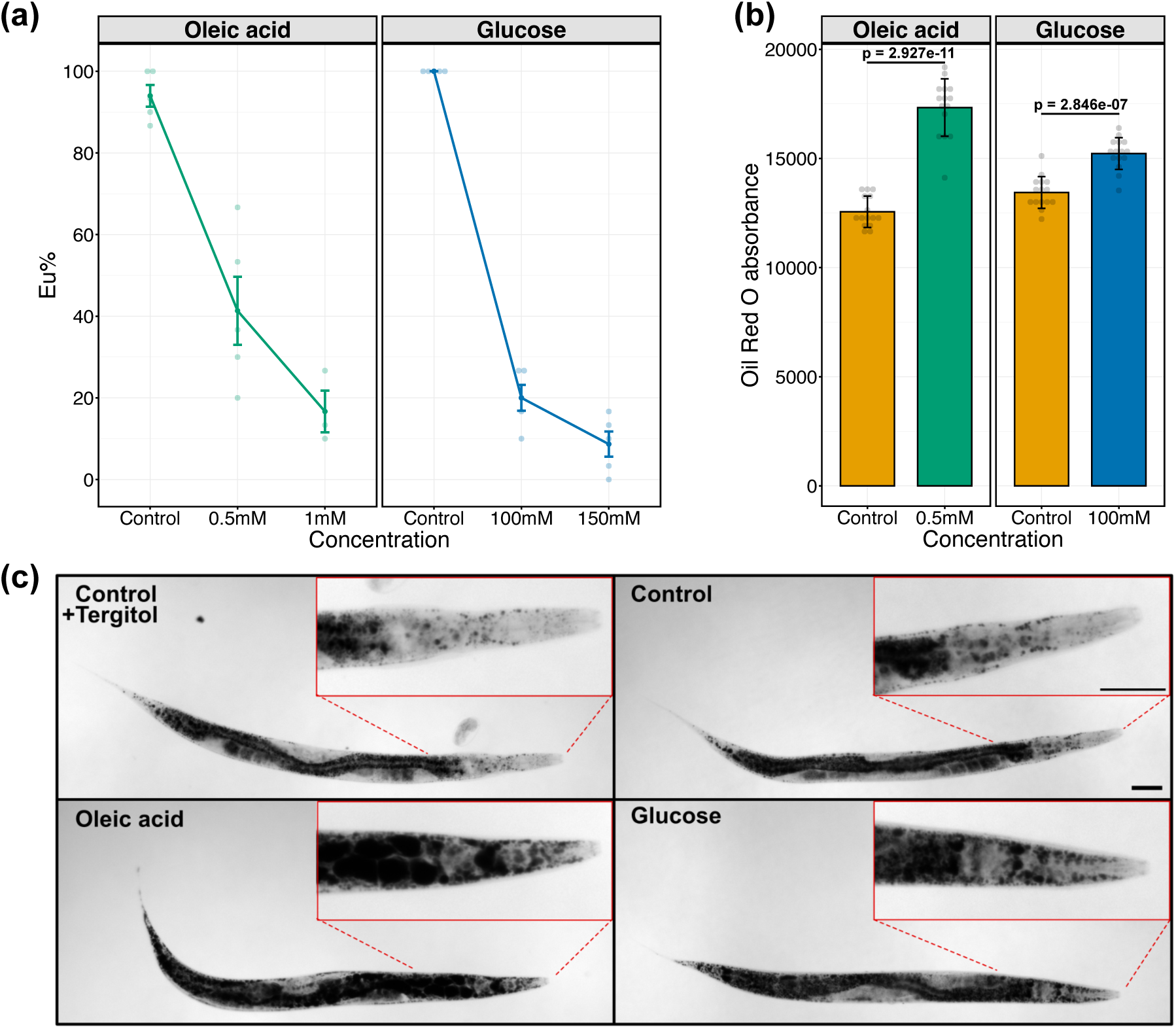
Effect of oleic acid and glucose supplementations on mouth-form plasticity and lipid storage in *P. pacificus*. (a) Concentration-dependent effect of oleic acid and glucose on mouth-form plasticity. N ≥ 3 biological replicates per condition for each concentration. Each faint data point represents a replicate (plate), with 30 animals per replicate being scored for mouth-form percentage (Eu%). Error bars represent s.e.m. (b) Oil Red O absorbance (RawIntDen/body area) obtained from worms grown in oleic acid, glucose, and respective control conditions. N = 15 per condition. Each faint data point represents an individual worm. P values are obtained from Welch two sample t-test and two sample t-test for oleic acid and glucose comparisons, respectively. Bars represent mean values of all samples in each condition. Error bars represent s.d. (c) Representative images of ORO-quantified worms, indicating lipid storage profile. Images are acquired from the blue channel in grayscale. Lipid droplets appear dark. Scale bar is 50μm.

### Oleic acid and glucose diets do not induce a transgenerational effect on mouth-form plasticity

Some dietary and nutritional influences on nematodes are known to be transmitted transgenerationally (Beltran et al., 2020). Therefore, we propagated cultures on oleic acid and glucose conditions for five generations to explore potential transgenerational effects of these diets on mouth-form plasticity (Figure S2a,b). We found that worms exhibit similar mouth-form ratios relative to their initial response (F1) through generations (Figure S2c,d). When we reverted worms from each generation back to the regular dietary condition without supplements, we did not observe any transgenerational memory of mouth form (Figure S2e,f). Note that over the course of these experiments, we observed more consistent and less variable mouth-form responses in glucose condition (highly St, on average below 20% Eu) relative to oleic acid across different batches.

### Oleic acid and glucose supplementations induce fat storage

Next, we performed ORO staining on worm populations reared on oleic acid and glucose diets to visualise and quantify their lipid storage (Figure 1b). Relative to control groups, oleic acid- and glucose-fed worms showed an increase in ORO absorbance which signifies an increase in fat storage (Figure 2b). Whole-body images of stained animals also demonstrated the distribution of storage lipids. Anterior regions of worms clearly indicated lipid accumulation in both oleic acid and glucose conditions (Figure 2c). In particular the head region of the worm allows easy visualization of the influence of supplementation on fat storage (Figure 2c inlets). Thus, both tested supplements induce fat storage in addition to their effect on mouth-form expression.

### Differential gene expression analysis points out pathways involved in lipogenesis and lipolysis

To understand which molecular factors are involved in mediating this nutrition-induced mouth-form change, we carried out RNA-seq on worm populations reared in supplemented and control conditions (Figure 3a). First, we performed differential gene expression analysis between each supplement and its control, and then a KEGG enrichment analysis of differentially expressed genes in both oleic acid and glucose conditions. KEGG enrichment analysis revealed pathways involved in lipogenesis (“biosynthesis of unsaturated fatty acids”, “fatty acid elongation”) and lipolysis (“peroxisome”, “fatty acid degradation”) (Figure 3b). This suggested that worms do not only store but also actively utilise lipids in supplemented conditions. Furthermore, we found genes which were upregulated and involved in peroxisomal beta-oxidation and biosynthesis of unsaturated fatty acids. Therefore, we selected genes in these pathways as candidates to investigate whether they mediate the effect of nutrient supplementation on mouth-form expression. We utilised CRISPR gene editing technology to introduce mutations in genetic components involved in both pathways as this method allows easy manipulation in *P. pacificus* (Han et al., 2020; Witte et al., 2015). For these experiments, we only used glucose-supplemented diet to induce fat storage and test responses of mutants because of the higher reproducibility of results obtained through glucose supplementation and also the impracticality of fatty acid supplementations that require preventive measures from oxidation by light and air (see Materials and methods for more details).

**Figure 3.**
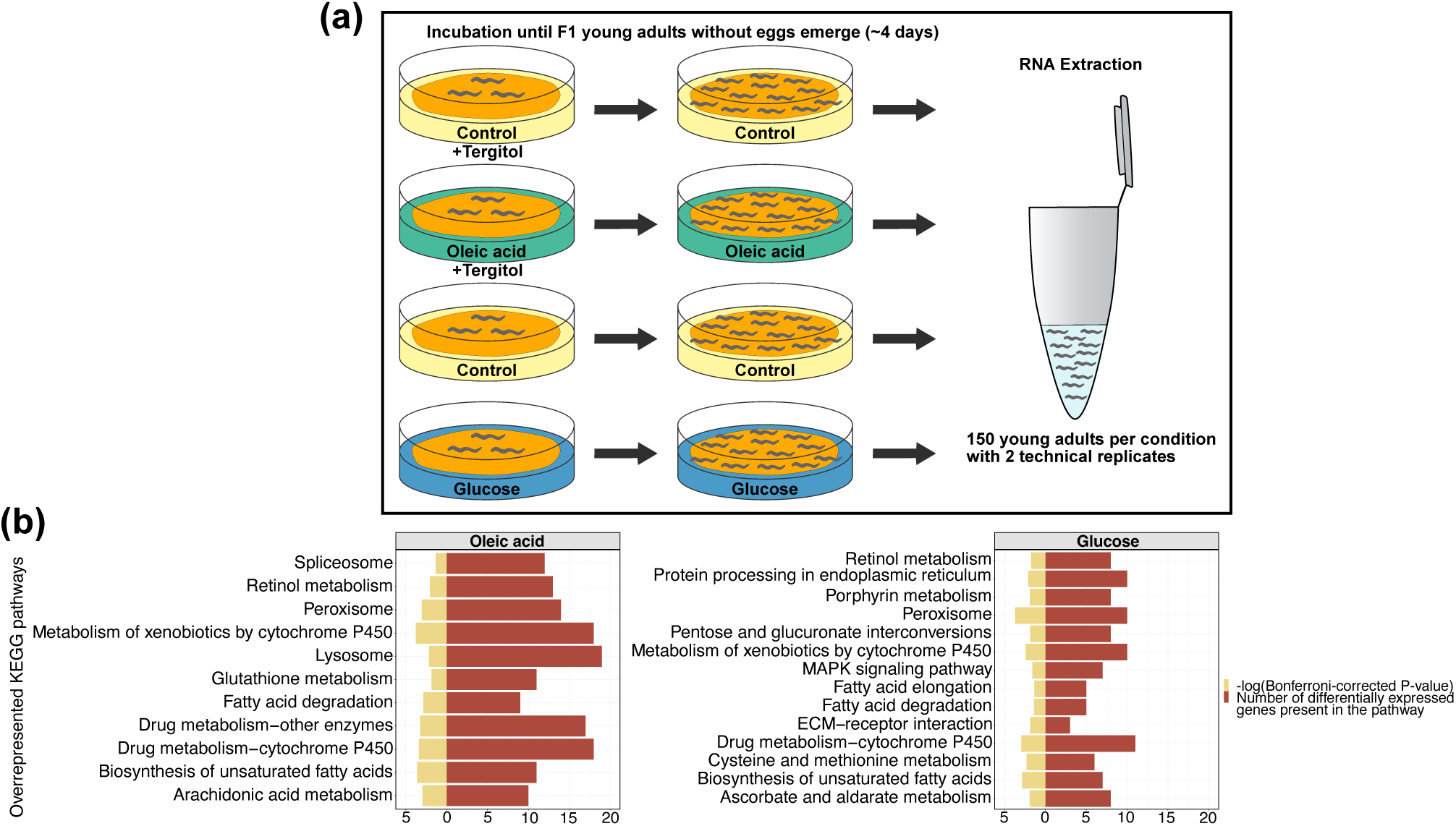
Transcriptome analysis. (a) An illustration of RNA extraction. (b) Overrepresented KEGG pathways of differentially expressed genes for oleic acid and glucose conditions. Statistical analysis by Fisher’s exact-test with multiple testing correction (Bonferroni corrected P<0.05).

### Delta-9 desaturase activity is essential for a complete mouth-form response on glucose-supplemented diet

We targeted delta-9 fatty acid desaturases for studying the role of the biosynthesis of unsaturated fatty acids and *dhs-28* and *daf-22* for investigating the peroxisomal beta-oxidation pathway on nutritional plasticity of the *P. pacificus* mouth-form. Delta-9 desaturases facilitate lipid storage by acting early in the *de novo* fatty acid synthesis pathway, converting saturated fatty acids into monounsaturated fatty acids (Figure 4a). In *C. elegans,* for instance, simultaneous inhibition of the function of two delta-9 desaturases (*fat-6* and *fat-7*) results in the loss of endogenous unsaturated fatty acids and a significant reduction in lipid storage (Brock et al., 2007). Delta-9 desaturases are activated by the transcription factor *sbp-1* (sterol regulatory element binding protein). Like *C. elegans*, *P. pacificus* has a single *sbp-1* gene (PPA37968), which is a one-to-one ortholog of *Cel-sbp-1*. First, we introduced frameshift mutations in this gene to block the activity of delta-9 desaturases and explore its contribution to mouth-form plasticity. However, homozygous mutants were sterile; therefore, we could not further our investigation with this gene. Next, we specifically targeted individual delta-9 desaturases. Interestingly, this gene family has been amplified in *P. pacificus* relative to *C. elegans* (Markov et al., 2015). We constructed a phylogenetic tree based on protein sequences of all *P. pacificus* fatty acid desaturase domain-containing genes and related *C. elegans* desaturases. We found a total of 10 *P. pacificus* genes which contain a delta-9 fatty acid desaturase-like protein domain (Figure S3). Note that unlike for *sbp-1*, there is no one-to-one orthology relationship between delta-9 fatty acid desaturase genes between *P. pacificus* and *C. elegans*, therefore requiring a new nomenclature. Among these genes, ppa_stranded_DN27845_c0_g3_i3, PPA1053, PPA40514, and ppa_stranded_DN27845_c0_g2_i1 showed highest protein sequence similarity with *C. elegans* delta-9 desaturases (*fat-5*, *fat-6*, and *fat-7*). In this order, we named these genes as *Ppa-pddl-1* (*Pristionchus* delta-9 desaturase-like-1), *Ppa-pddl-2*, *Ppa-pddl-*3, and *Ppa-pddl-4* (Figure 4b, Figure S3). Among all, *Ppa-pddl-1*, *Ppa-pddl-3*, and *Ppa-pddl-4* are expressed throughout the development of *P. pacificus* based on previous gene expression analysis (Baskaran et al., 2015). Therefore, we prioritised these three genes and introduced mutations through CRISPR. We obtained three frameshift alleles for both *Ppa-pddl-3* (*tu2033*, *tu2034*, *tu2035*) and *Ppa-pddl-4* (*tu2030*, *tu2031*, *tu2032*), and one frame shift allele (*tu2028*) and a 3bp insertion allele (*tu2029*) for *Ppa-pddl-1* (Table S2). Among these mutants, *tu2028* and *tu2029* were developmentally slow and had significantly reduced fat storage until late adulthood under standard conditions.

**Figure 4.**
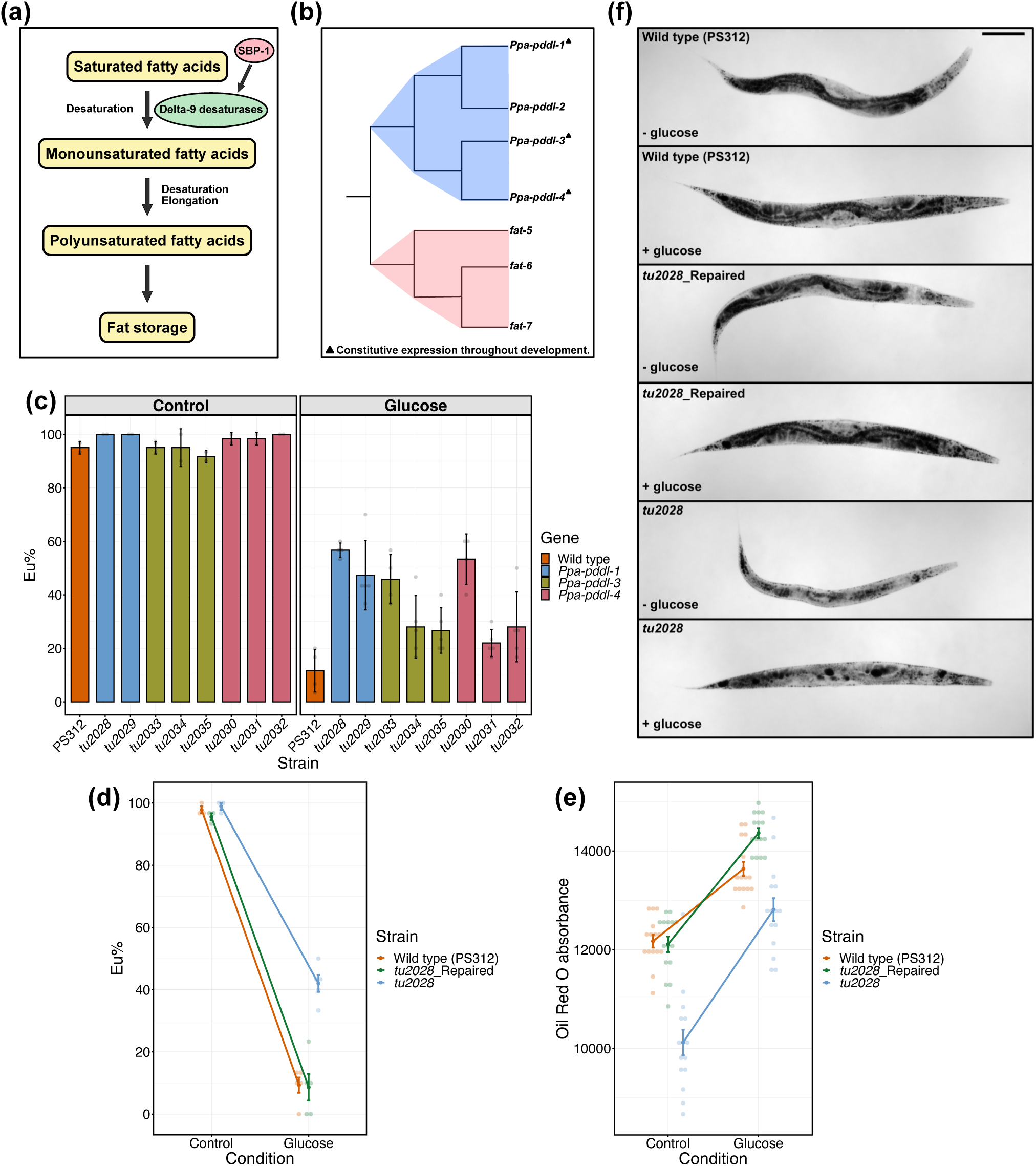
Significance of delta-9 desaturase activity in glucose supplementation-induced mouth-form plasticity. (a) Schematic representation of the essential activity of delta-9 desaturases, facilitating fat storage by converting saturated fatty acids into monounsaturated fatty acids. Sterol regulatory element binding protein 1 (SBP-1) activates delta-9 desaturases. (b) A simplified phylogenetic tree of *Pristionchus* delta-9 desaturase-like genes (*Ppa-pddl-1*, *Ppa-pddl-2*, *Ppa-pddl-3*, *Ppa-pddl-4*), and *C. elegans* delta-9 desaturases (*fat-5*, *fat-6*, and *fat-7*). Triangle denotes *P. pacificus* genes which are constitutively expressed during development (Baskaran et al., 2015). The phylogenetic tree is drawn based on the tree in Figure S3. (c) Eu percentages of CRISPR mutants of *Ppa-pddl-1*, *Ppa-pddl-3*, *Ppa-pddl-4*, and wild type animals in control and glucose conditions. N ≥ 2 biological replicates per strain for each condition. Bars represent mean values of all replicates. Error bars represent s.d. (d) Eu percentages of wild type, *tu2028*(*Ppa-pddl-1*), and CRISPR repaired-*tu2028* strains in control and glucose conditions. N ≥ 3 biological replicates per strain for each condition. (c,d) Each faint data point represents a replicate (plate), with 30 animals per plate being scored for mouth-form percentage (Eu%). (e) ORO absorbance (RawIntDen/body area) obtained from wild type, *tu2028*(*Ppa-pddl-1*), and CRISPR repaired-*tu2028* strains grown on control and glucose conditions. N = 15 per strain for each condition. Each faint datapoint represents an individual worm. (d,e) Error bars represent s.e.m. (f) Representative images of ORO-quantified strains from control (-glucose) and glucose (+glucose) conditions. Images are acquired from the blue channel in grayscale. Lipid droplets appear dark. Scale bar is 100μm.

Next, we assessed mouth-form responses of all mutants on glucose-supplemented diets. Results revealed that the response of delta-9 desaturase mutants to glucose is generally incomplete, that is, they have a higher Eu ratio than wild type animals (above 20% Eu) (Figure 4c). Compared to all other mutants, *Ppa-pddl-1* mutants were more consistent in their mouth-form response to the glucose-supplemented diet (greater than 40% Eu on average) (Figure 4c). Therefore, we further studied *Ppa-pddl-1* using *tu2028* as reference allele. First, we measured the lipid storage profile of *Ppa-pddl-1*(*tu2028)* on standard diet by ORO staining. We found that *Ppa-pddl-1*(*tu2028)* exhibits significantly lower levels of ORO absorbance relative to wild type (Figure S4a,b). To confirm that the response of *tu2028* to glucose-supplemented diet is due to its mutational background in *Ppa-pddl-1* and not to other mutations as a result of potential off target effects by CRISPR, we genetically reverted *Ppa-pddl-1*(*tu2028)* back to wild type using CRISPR editing via a repair template. Growing wild type, the repaired strain, and the original *Ppa-pddl-1*(*tu2028)* mutant on glucose-supplemented diets revealed highly similar levels of mouth-form ratios between wild type and repaired worms (Figure 4d). Specifically, both strains were highly St on glucose-supplemented diets. Repaired and wild type strains also showed a similar trajectory of ORO absorbance between control and glucose conditions (Figure 4e). Among all, *Ppa-pddl-1*(*tu2028)* had the weakest ORO absorbance in both conditions (Figure 4e). In addition, we observed that *Ppa-pddl-1*(*tu2028)* mutant animals restored their growth on glucose-supplemented diet and also regained their wild type physiological and morphological characteristics upon genetic repair (Figure 4f). Overall, these results highlight the significance of the function of *Ppa-pddl-1* in both lipid storage and mouth-form plasticity.

### Peroxisomal beta-oxidation mutants fail to respond to glucose supplementation

Next, we studied mutants of two peroxisomal beta-oxidation genes, *dhs-28* and *daf-22*. *P. pacificus* has two copies of both genes in its genome (Markov et al., 2016) (Figure 5a). For *Ppa-dhs-28.1*, we induced mutation in the copy with the sterol carrier protein domain (PPA20393) and obtained two insertion and three deletion alleles (Table S2). All mutants exhibited the same observable characteristics and were morphologically smaller and developmentally slower relative to wild type animals. For *daf-22*, we used the available double mutant *Ppa-daf-22.1 Ppa-daf-22.2* (Markov et al., 2016). Note that mutants of *Ppa-dhs-28.1* were generated in the PS312 wild type background, whereas the *Ppa-daf-22.1 Ppa-daf-22.2* double mutant was generated in RS2333, which is a highly related derivative of PS312 that was used for several studies on dauer development (Falcke et al., 2018; Markov et al., 2016). Therefore, we included both highly similar wild type strains in our experiments as independent controls. First, we assessed mouth-form responses of three *Ppa-dhs-28.1* mutant alleles (*tu1855*, *tu1856*, and *tu1858*) and the *Ppa-daf-22.1 Ppa-daf-22.2* double mutant on glucose-supplemented diets. Mutants of both genes had a consistently high Eu mouth-form ratio between control and glucose conditions. Next, we simultaneously assessed mouth-form plasticity and lipid storage of *Ppa-dhs-28.1(tu1855*) and the *Ppa-daf-22.1 Ppa-daf-22.2* double mutant on glucose-supplemented diets. Relative to both wild type strains, mutants remained highly predatory and showed no increase in ORO absorbance between control and glucose conditions (Figure 5b,c). We also observed that glucose-supplemented peroxisomal beta-oxidation mutants exhibit delayed development and reduced body size. Moreover, ORO-stained peroxisomal beta-oxidation mutants show a disruption in lipid storage integrity on glucose-supplemented diets (Figure 5d). They also accumulate larger lipid droplets relative to wild type, indicating reduced lipid oxidation. Taken together, these results indicate that the peroxisomal beta-oxidation pathway is required for the mouth-form response to glucose.

**Figure 5.**
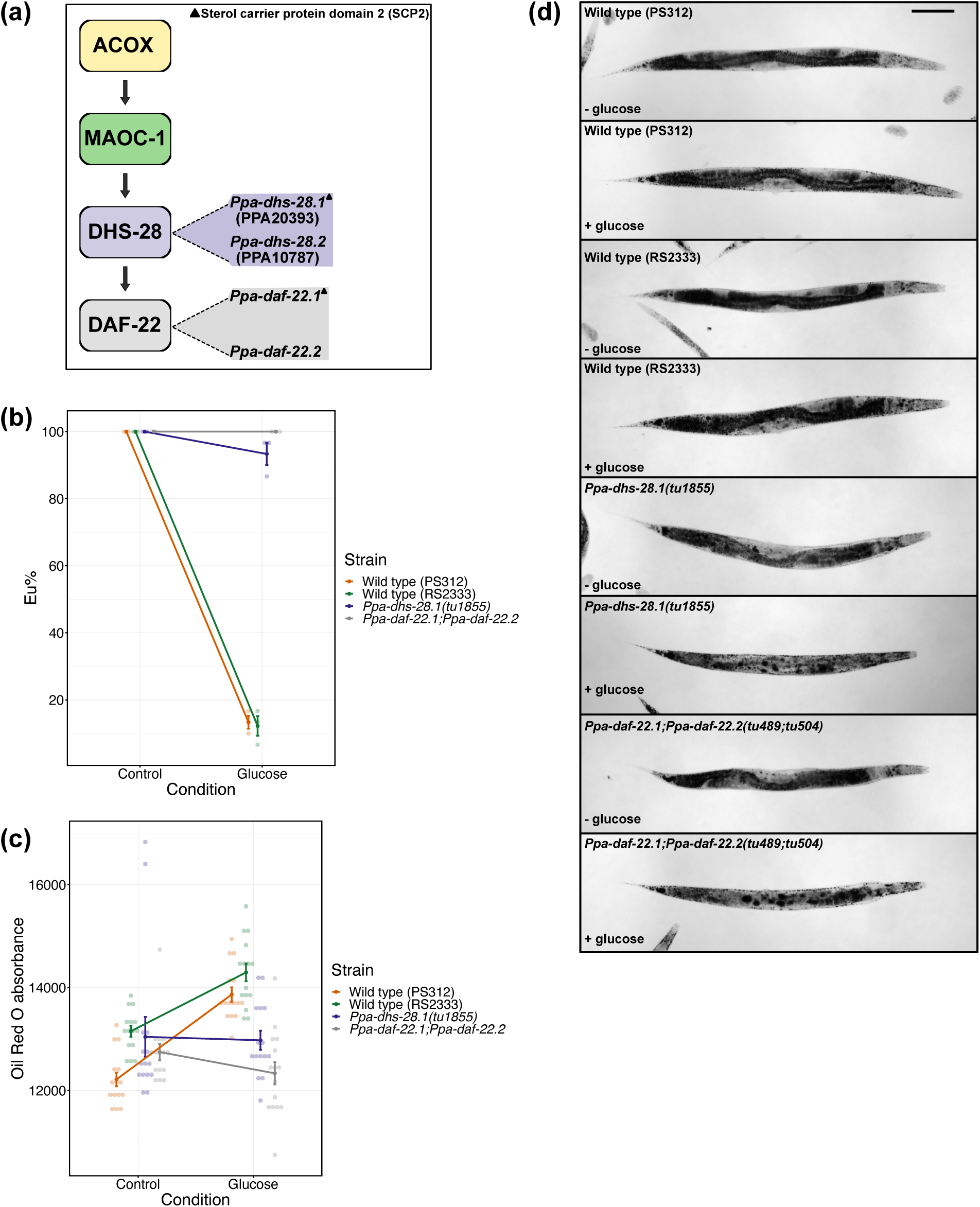
Response of peroxisomal beta-oxidation mutants to glucose-supplemented diet. (a) Schematic of peroxisomal beta-oxidation pathway, indicating duplicate copies in *P. pacificus* for *dhs-28* and *daf-22* genes relative to *C. elegans*. (b) Eu percentages of peroxisomal beta-oxidation mutants and wild type strains in control and glucose conditions. N ≥ 2 biological replicate per strain for each condition. Each faint data point represents a replicate (plate), with 30 animals per plate being scored for mouth-form percentage (Eu%). (c) ORO absorbance (RawIntDen/body area) obtained from peroxisomal beta-oxidation mutants and wild type strains grown on control and glucose conditions. N = 15 per strain for each condition. Each faint data point represents an individual worm. (b,c) Error bars represent s.e.m. (d) Representative images of ORO-quantified strains from control (-glucose) and glucose (+glucose) conditions. Images are acquired from the blue channel in grayscale. Lipid droplets appear dark. Scale bar is 100μm.

### Mouth-form plasticity switch genes are required for phenotypic response in glucose diet

Next, we tested whether the influence of glucose-supplemented diets on mouth-form plasticity is mediated through the central gene regulatory network of mouth-form plasticity (Figure 6a). For that, we utilised mutants of plasticity switch genes which are St-form defective. Specifically, we used *sult-1(tu1061)* and the double mutant *nag-1(tu1142) nag-2(tu1143*) and assessed their mouth form and lipid storage responses on glucose-supplemented diets. Both mutants were unable to switch from Eu to St on glucose-supplemented diets, indicating that the nutritional effect acts upstream of the plasticity switch module (Figure 6b). However, plasticity switch mutants were able to increase their lipid storage when fed on glucose-supplemented diets (Figure 6c). This suggests that the plasticity switch genes are essential for mouth-form response but not for lipid storage.

**Figure 6.**
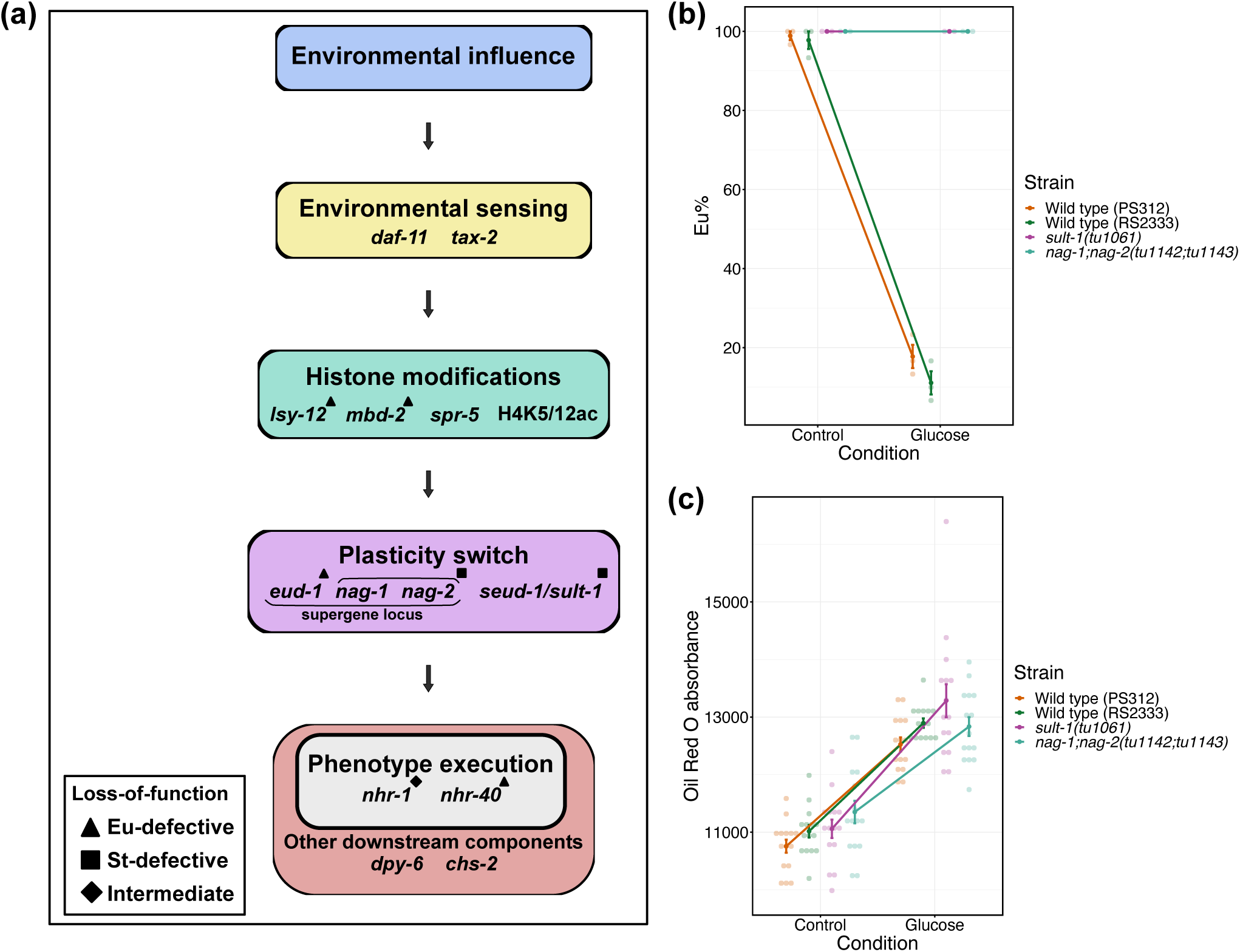
Role of plasticity switch genes in glucose supplementation-induced mouth-form plasticity. (a) Schematic of the mouth-form gene regulatory network, indicating different modules with genetic and epigenetic components. (b) Eu percentages of plasticity switch mutants and wild type strains in glucose and control conditions. N = 3 biological replicates per strain for each condition. Each faint data point represents a replicate (plate), with 30 animals per plate being scored for mouth-form percentage (Eu%). (c) ORO absorbance (RawIntDen/body area) measured in plasticity switch mutants and wild type strains grown on control and glucose conditions. N = 15 per strain for each condition. Each faint data point represents a worm. (b,c) Error bars represent s.e.m.

### Non-predatory worms exhibit a fitness advantage over predatory ones

Our final aim was to explore whether there is a fitness advantage of favouring the development of one morph over the other in a fat storage-inducing condition. For this, we chose fecundity as proxy for fitness to evaluate the hermaphrodites’ reproductive success by counting viable progeny produced by isolated individuals. First, we reduced the concentration of glucose from 100mM to 80mM to increase the likelihood of obtaining Eu animals. Then, we isolated animals of both morphs grown in the glucose-supplemented diets and measured their fecundity on the standard diet (Figure 7a). Results revealed that St worms on average exhibit higher daily and total fecundity than Eu worms (Figure 7b,c). These findings suggest that promoting the development of the St morph under such a dietary influence confers a fitness advantage to *P. pacificus*.

**Figure 7.**
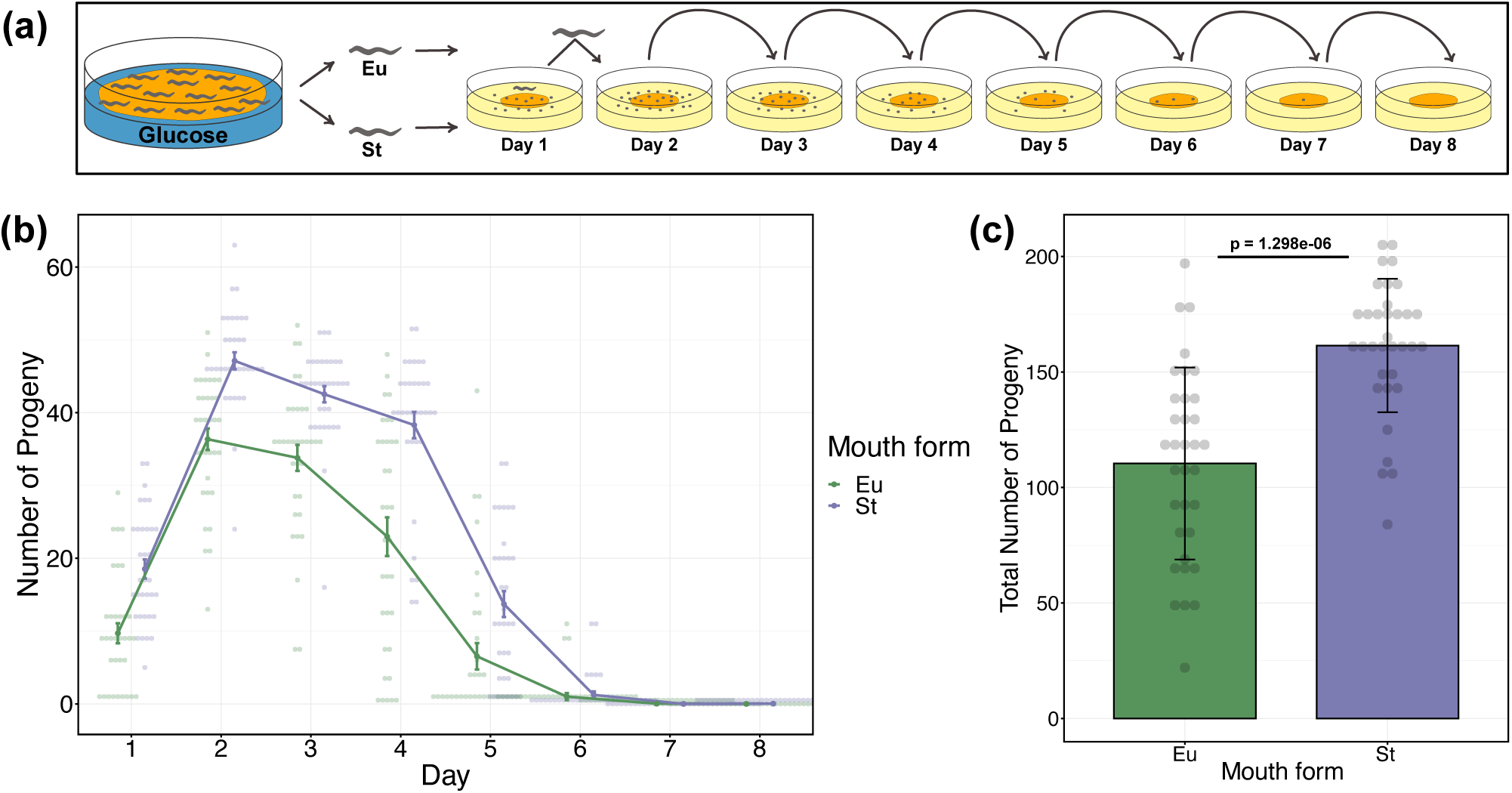
Fecundity of different morphs after glucose supplementation. (a) Illustration for the experimental design to study fecundity of Eu and St animals obtained from glucose-supplemented condition. (b) Daily fecundity of Eu and St animals. Error bars represent s.e.m. (c) Overall fecundity of Eu and St animals. Bars represent mean values of all individuals for each morph. Error bars represent s.d. P value is obtained from Wilcoxon rank sum test. (b,c) N = 34 per morph. Each faint data point represents a worm.

## Discussion

This study highlights the significance of nutrition in mouth-form plasticity in *P. pacificus*. First, we showed that fat storage-promoting conditions induce the non-predatory morph. This suggests that growing in an “overly satiated” nutritional state increases the likelihood of worms developing as non-predatory. These findings are consistent with previous studies, which indicated that opposite dietary condition, *i.e.*, low nutrition, can lead to the predatory morph in *P. pacificus* (Bento et al., 2010); and even a cannibalistic novel predatory mouth-form in another diplogastrid nematode, *Allodiplogaster sudhausi* (Wighard et al., 2024). In addition, results obtained from transgenerational experiments suggest that worms must constantly be kept at overly satiated state during development to obtain and maintain a mouth-form response, at least in the wild type *P. pacificus* PS312 strain.

Supplementing NGM agar plates with glucose and oleic acids has been an effective method for delivering monosaccharides and fatty acids to worms via dietary ingestion (Alcántar-Fernández et al., 2018; Deline et al., 2013; Nomura et al., 2010). However, it is important to note that this method does not address any potential indirect effect caused by the interaction between the supplement and the bacteria (Kingsley et al., 2021), which can be a research interest on its own. In our experiments, we did not monitor fatty acid and glucose uptake or content in worms, instead we utilised ORO staining, to monitor the outcome of changes in lipid storage. Our results are concordant with previous findings. For instance, glucose supplementation to NGM plates does not only result in the accumulation of glucose but also triacylglycerols in *C. elegans* (Alcántar-Fernández et al., 2018). Oil Red O stains neutral lipids, mainly triacylglycerols. Studies in *C. elegans* have shown an increase in ORO levels in worms grown in oleic acid- and glucose-supplemented conditions (Choi et al., 2021; Lee et al., 2015). Therefore, ORO does not only provide a way of measuring the nutritional status of worms but also validate the uptake of supplementations. Moreover, our results showed that glucose supplementation consistently induces the St morph in higher proportions relative to oleic acid. Note that we could not rule out the cause of inconsistency and the high variability of mouth-form ratios in oleic acid supplementations. Regardless, we selected glucose-supplemented diet as the main fat storage-promoting condition to induce the non-predatory morph.

Further, mutant analyses revealed that genes involved in lipogenesis and lipolysis play a significant role in glucose supplementation-induced mouth-form plasticity. First, we showed that delta-9 desaturase activity, particularly *Ppa-pddl-1*, is required for a complete mouth-form and lipid storage response. The spatial transcriptome data, published by Rödelsperger et al. (2021), indicates that *Ppa-pddl-1* is enriched in anatomical regions that suggests expression in the intestine. Inactivation of this gene causes a drastic reduction in lipid storage and developmental rate; such phenotypic effect is obtained through simultaneous inhibition of *fat-6* and *fat-7* in *C. elegans* (Brock et al., 2007). However, *Ppa-pddl-1* mutant still exhibits a partial ability to respond to glucose-supplemented diet. This suggests that remaining functional delta-9 desaturases may have compensated for the loss of *Ppa-pddl-1*. Therefore, further functional characterization of delta-9 desaturases is required to completely elucidate potential role of this gene family in mouth-form plasticity. Furthermore, growing peroxisomal beta-oxidation mutants on glucose-supplemented diet revealed that these mutants have disrupted lipid storage integrity, which resulted in a weaker ORO absorbance relative to wild type strains. Additional observations in these mutants, such as developmental delay and reduction in body size, suggest that they are not able to nutritionally benefit from the glucose-supplemented diet. Taken together, these findings suggest that pathways associated with storage and utilisation of lipids carry out essential metabolic processes, mediating mouth-form plasticity in a high nutrition environment.

Additionally, our findings suggest that the dietary effect of glucose supplementation acts upstream of the plasticity switch module. However, how this dietary effect is mediated to influence downstream components, affecting mouth-form decision remains subject of future research. Of note, *de novo* fatty acid synthesis and peroxisomal beta-oxidation pathways produce signalling molecules which have diverse functions (Artyukhin et al., 2018; Watts, 2016). For example, polyunsaturated fatty acids produced by the *de novo* fatty acid synthesis pathway can be integrated into storage lipids or processed further into signalling molecules such as eicosanoids, which can function as ligands for transcription factors, affecting gene expression. Cytochrome P450 enzymes are involved in the production of eicosanoids from polyunsaturated fatty acids (Kulas et al., 2008) and their activity have been associated with several biological functions such as development, dauer formation, and lipid metabolism in *C. elegans* (for review, see Larigot et al., 2022). Cytochrome P450-related pathways are enriched (KEGG) in our differential gene expression data for both oleic acid and glucose, suggesting a potential role. Moreover, peroxisomal beta-oxidation pathway produces ascarosides, extracellular signalling molecules which can influence dauer formation and mouth-form plasticity (Butcher, 2017; Butcher et al., 2007; Markov et al., 2016; Werner et al., 2018). Taken together, lipid-mediated signalling may have potential functions in nutrition-induced mouth-form plasticity and therefore further research is required to elucidate associated mechanisms.

Finally, we sought to explore whether there is an adaptive value of facilitating the development of the St morph in a high nutrition environment. In *P. pacificus*, while the Eu morph exhibits a fitness advantage associated with its predatory ability (Serobyan et al., 2014), the St morph claims this advantage through a faster development and a higher fecundity (Dardiry et al., 2023; Serobyan et al., 2013). In addition, highly non-predatory natural isolates of *P. pacificus* have higher fecundity and exhibit faster development on standard dietary condition relative to highly predatory strains, which were isolated from similar localities in nature (Dardiry et al., 2023). Growing worms on glucose-supplemented diet allowed separation of morphs; then assessing their fecundity revealed a fitness disadvantage for the predatory mouth-form. Intriguingly, our findings suggest that glucose supplementation does not offset the fitness cost of producing the Eu phenotype or promote its development to begin with. Hence, the fitness advantage associated with the non-predatory morph suggests that favouring its development, under such dietary effect, can be beneficial for the population.

In summary, this study signifies the nutritional sensitivity of mouth-form polyphenism in *P. pacificus* and adds nutritional status as important environmental factor influencing mouth form. We introduced glucose-supplementation as a novel environmental condition to induce non-predatory morph. Our findings suggest that nutritional status of the worm can potentially dictate its mouth-form fate. We also found a strong association between lipid metabolism and mouth-form plasticity. Lastly, glucose-supplementation helped understand fitness consequences of the mouth-form determination, indicating an advantage for the non-predatory morph.

## Acknowledgements

We would like to thank Dr Christian Rödelsperger for his guidance and support during this project. We also thank Heike Haussmann for freezing all the worm strains generated by this work. Additionally, we thank all the current and past members of the SommerLab for valuable discussions.

## Data availability

Raw Reads obtained from the RNA-seq were deposited in the European Nucleotide Archive (ENA) under the accession number PRJEB76787.

