## Supplementary material for "High nutritional conditions influence feeding plasticity in *Pristionchus pacificus* and render worms non-predatory"

**Supplementary Figures and Tables**

**Veysi Piskobulu[1], Marina Athanasouli[1], Hanh Witte[1], Christian Feldhaus[2], Adrian Streit[1] & Ralf J. Sommer[1,*]**

[1] Max-Planck Institute for Biology Tübingen, Department for Integrative Evolutionary Biology, Max-Planck Ring 9, D-72076 Tübingen, Germany

[2] Max-Planck Institute for Biology Tübingen, BioOptics Facility, Max-Planck Ring 5, D-72076 Tübingen, Germany

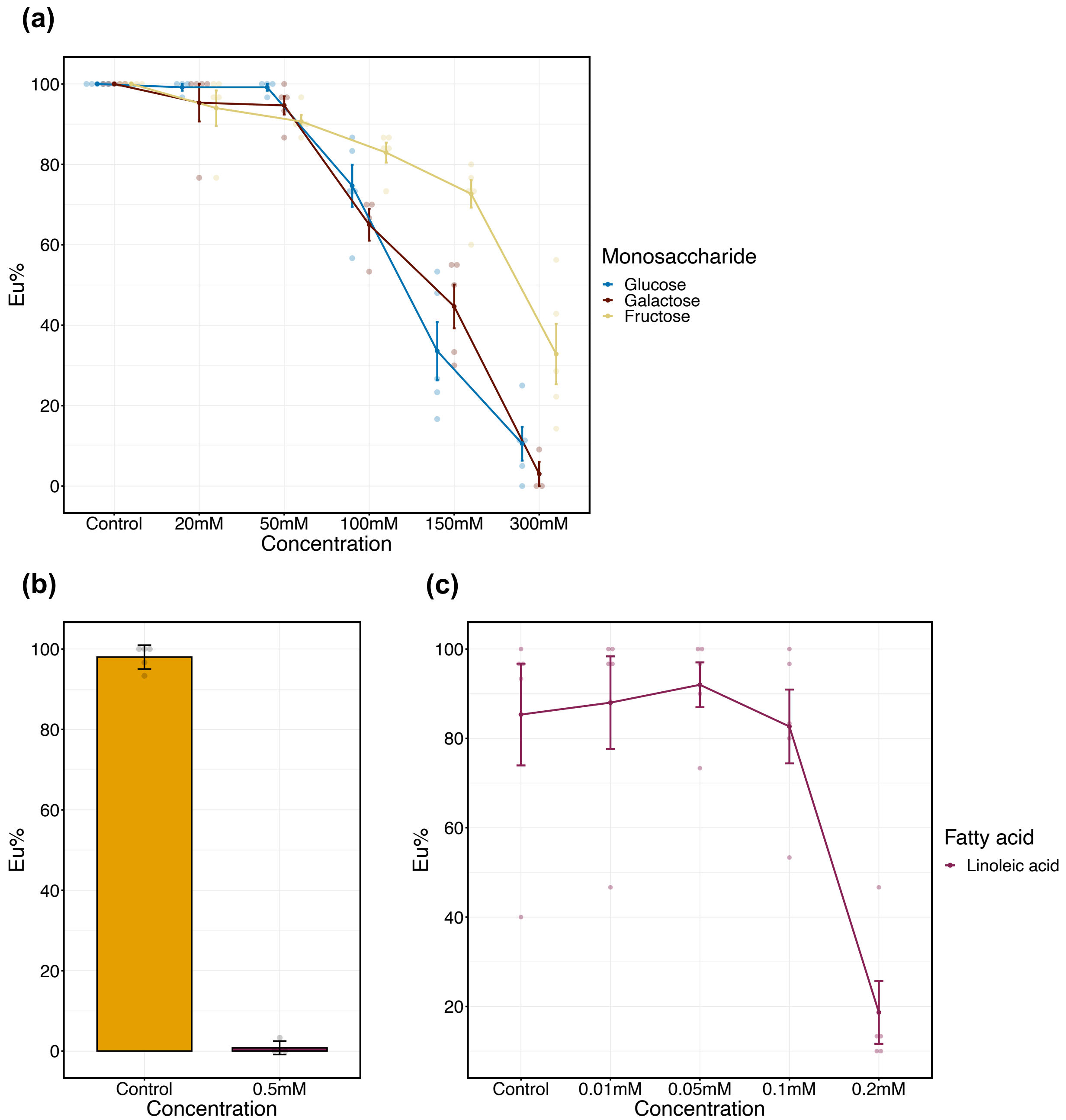


**Supplementary Figure S1**

**Concentration dependent effect of supplements on mouth-form plasticity.** (a) Eu percentages of worms grown on different concentrations of monosaccharides. In this pilot experiment, cultures were initiated by inoculating with 10 adult worms, allowing them to lay eggs for 2 hours. This method results in higher Eu percentages than inoculating with 3 worms. N ≥ 3 biological replicates per condition in each concentration. From each replicate (plate), 25-30 worms were scored, except for 300mM concentrations from which 4-28 worms were scored. Error bars represent s.e.m. (b) Effect of 0.5mM linoleic acid on mouth-form plasticity. N ≥ 4 biological replicates per concentration. From each replicate (plate), 28-30 worms were scored. Bars represent mean values of all replicates. Error bars represent s.d. (c) Linoleic acid concentration effect on mouth-form plasticity. N = 5 biological replicates per concentration. From each replicate (plate), 30 worms were scored. Error bars represent s.e.m. (a-c) Each faint data point represents a biological replicate (plate) scored for mouth-form ratio (Eu%).

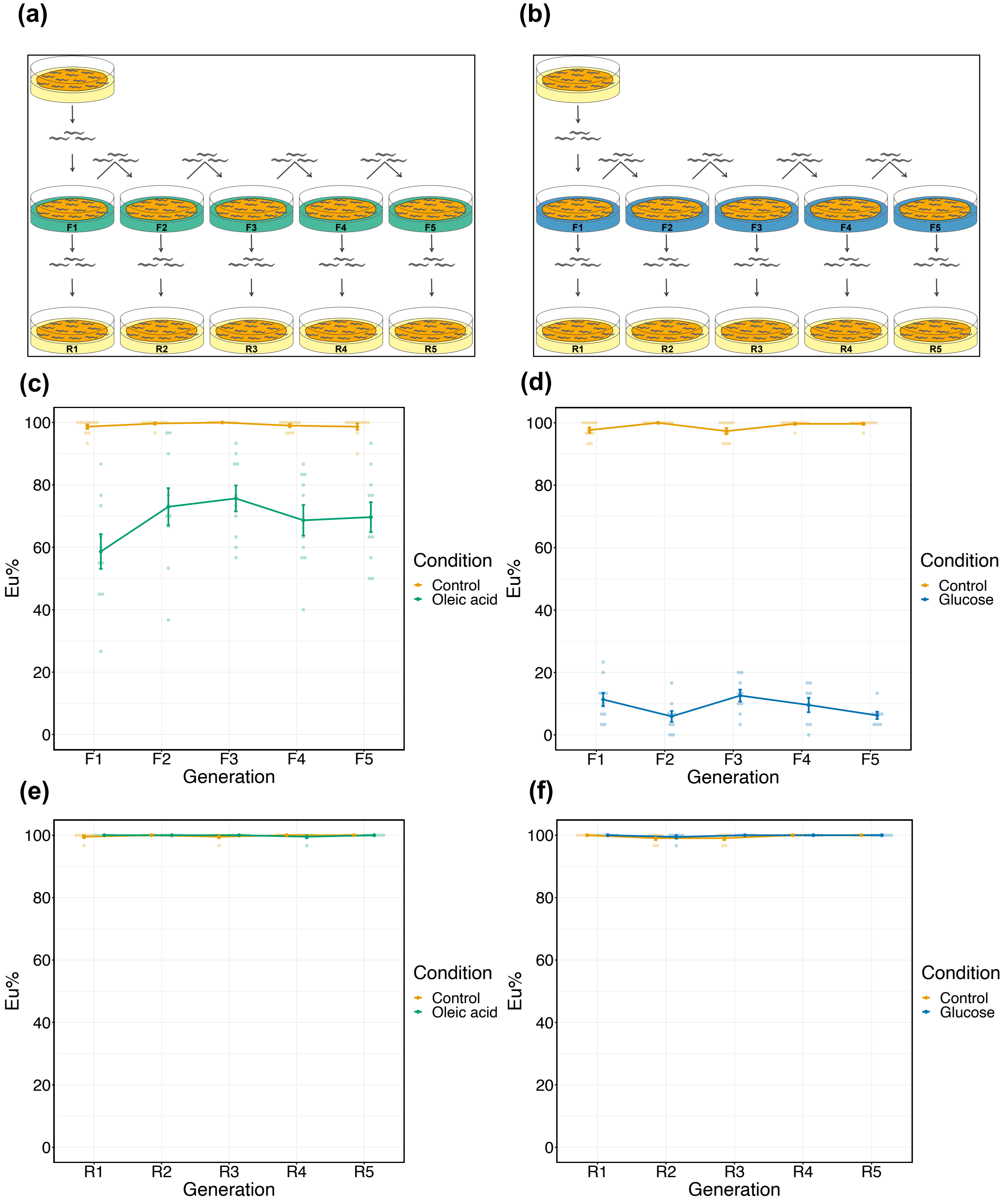


**Supplementary Figure S2**

**Transgenerational effect of oleic acid and glucose supplementations on mouth-form plasticity.** (a,b) Illustrations show the experimental design for studying the transgenerational effect of oleic acid (a) and glucose (b) supplementation on mouth-form plasticity. (a,b) The same experimental method was applied for respective control conditions. (c,d) Eu percentages of worms through generations (F1-F5) for oleic acid and control conditions (c); and for glucose and control conditions (d). (c) N = 10 biological replicates per condition for each generation. (d) N ≥ 8 biological replicates per condition for each generation. (e,f) Eu percentage of worms after reversal (F1-F5) for oleic acid and control conditions (e); and for glucose and control conditions (f). (e) N = 7 biological replicates per condition for each generation. (f) N ≥ 6 biological replicates per condition for each generation. (c-f) Each faint data point represents a replicate (plate), with 30 animals per plate being scored for mouth-form percentage (Eu%). Error bars represent s.e.m.

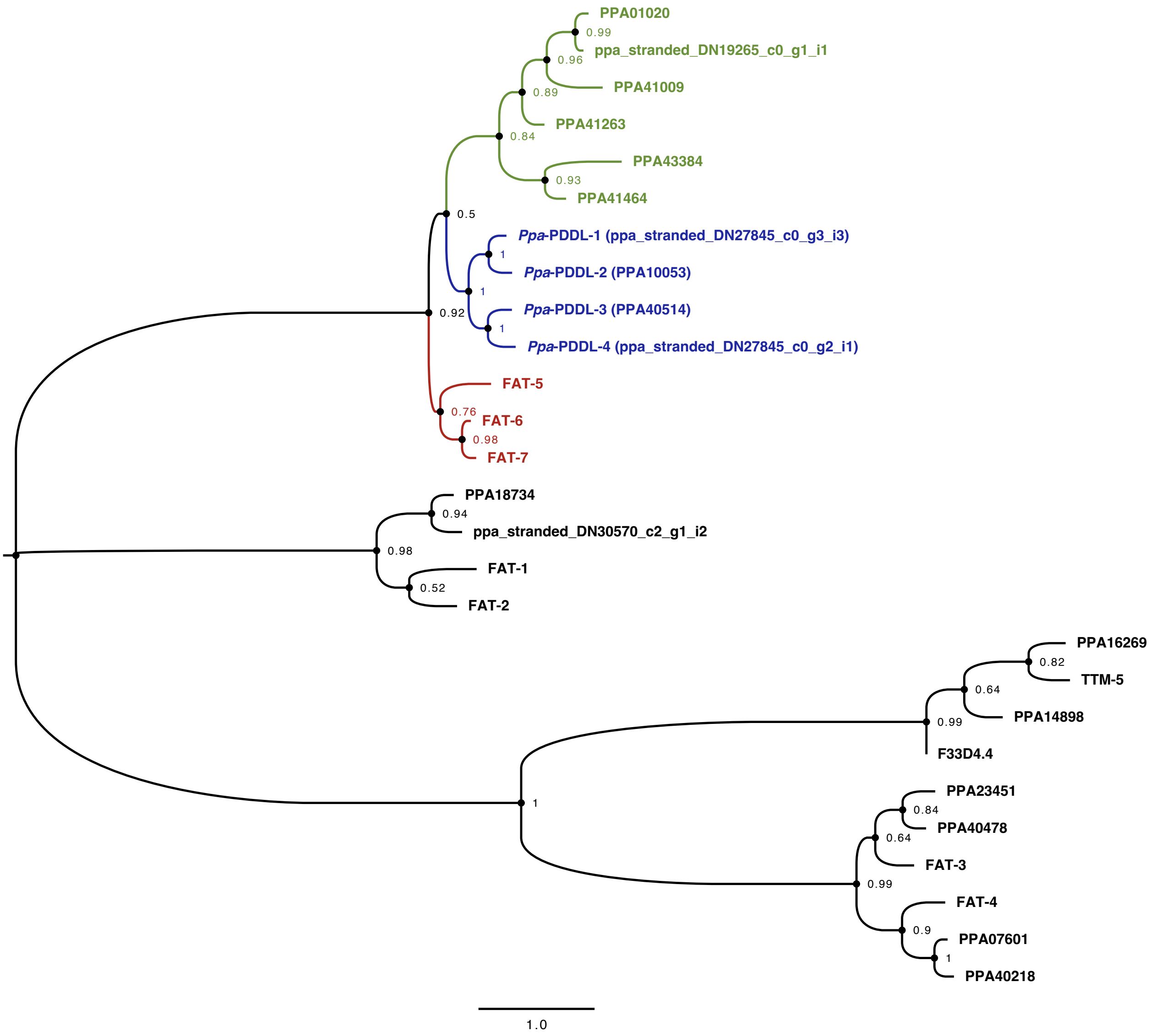


**Supplementary Figure S3**

**Phylogenetic tree of all *P. pacificus* fatty acid desaturase domain-containing proteins with related *C. elegans* desaturases.** A maximum likelihood phylogenetic tree, constructed with LG model, and 100 bootstrap replications. Bootstrap values are indicated next to branch nodes. Green and Blue colours denote *Pristionchus* delta-9 desaturase domain-containing proteins. Red colour denotes *C. elegans* delta-9 desaturases.

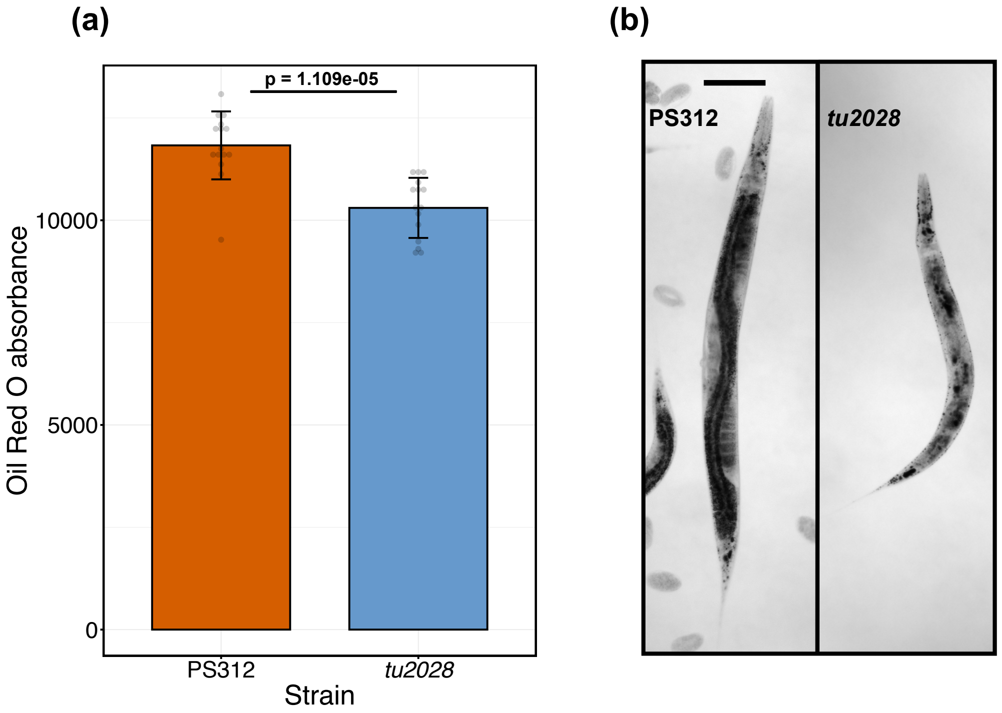


**Supplementary Figure S4**

**Delta-9 desaturase mutant, *Ppa-pddl-1(tu2028)*, exhibits reduced lipid storage relative to wild type strain (PS312) on standard dietary condition.** (a) ORO absorbance (RawIntDen/body area) obtained from wildtype (PS312) and *Ppa-pddl-1(tu2028)* strains. N = 15 per strain. P value is obtained from a two sample t-test. Each faint data point represents a worm. Bars represent mean values of all samples for each strain. Error bars represent s.d. (b) Representative images of ORO-quantified worms, indicating lipid storage profile. Images are acquired from the blue channel in grayscale. Lipid droplets appear dark. Scale bar is 100µm.

**Supplementary Table S1. List of crRNA and primer sequences for genes that were studied for mutant analyses.**

| Gene name | Accession | crRNA | Forward Primer | Reverse Primer |
|---|---|---|---|---|
| *Ppa-pddl-1* | ppa_stranded_DN27845_c0_g3_i3 | 5′-CAGACATTACTATCCTCTAG-3′ | 5′-CCGTTAGAGTCTACTTCATGCTATGGAA-3′ | 5′-ATCAACCTGACCATATTTCAGTCTGACC-3′ |
| *Ppa-pddl-3* | PPA40514 | 5′-CTATCTCCCCCTTGCGACTC-3′ | 5′-TTCTGATCTGTGGAACGACCCG-3′ | 5′-CACGGATTCGACGGGAGTGATG-3′ |
| *Ppa-pddl-4* | ppa_stranded_DN27845_c0_g2_i1 | 5′-GTAATTCCCGTCTATTTCTG-3′ | 5′-TAACGATGTTTTCCTTCAGGTAATGGGC-3′ | 5′-CTGCTGCTTGTAGATTAGTCCATACAGA-3′ |
| *Ppa-dhs-28.1* | PPA20393 | 5′-GGGGAGATCAAGGCAGCCGG-3′ | 5′-CGATATTGTTGCAGTGAACGAC-3′ | 5′-CTTCTAGTTACATCAGCTGTCTCG-3′ |

### Supplementary Table S2. List of CRISPR mutants utilised in this study.

| Gene Name | Gene Accession | Strain | Allele | Mutation Type | Mutation Location | Source |
|---|---|---|---|---|---|---|
| *Ppa-pddl-1* | ppa_stranded_DN27845_c0_g3_i3 | RS4411 | *tu2028* | 7bp insertion | exon 4 | This paper |
| *Ppa-pddl-1* | ppa_stranded_DN27845_c0_g3_i3 | RS4412 | *tu2029* | 3bp insertion | exon 4 | This paper |
| *Ppa-pddl-3* | PPA40514 | RS4401 | *tu2033* | 5bp deletion | exon 5 | This paper |
| *Ppa-pddl-3* | PPA40514 | RS4402 | *tu2034* | 7bp deletion | exon 5 | This paper |
| *Ppa-pddl-3* | PPA40514 | RS4403 | *tu2035* | 7bp deletion | exon 5 | This paper |
| *Ppa-pddl-4* | ppa_stranded_DN27845_c0_g2_i1 | RS4398 | *tu2030* | 4bp deletion | exon 7 | This paper |
| *Ppa-pddl-4* | ppa_stranded_DN27845_c0_g2_i1 | RS4399 | *tu2031* | 10bp deletion | exon 7 | This paper |
| *Ppa-pddl-4* | ppa_stranded_DN27845_c0_g2_i1 | RS4400 | *tu2032* | 28bp insertion | exon 7 | This paper |
| *Ppa-dhs-28.1* | PPA20393 | RS4147 | *tu1855* | 4bp deletion | exon 3 | This paper |
| *Ppa-dhs-28.1* | PPA20393 | RS4138 | *tu1856* | 7bp deletion | exon 3 | This paper |
| *Ppa-dhs-28.1* | PPA20393 | RS4141 | *tu1857* | 8bp insertion | exon 3 | This paper |
| *Ppa-dhs-28.1* | PPA20393 | RS4140 | *tu1858* | 21bp deletion | exon 3 | This paper |
| *Ppa-dhs-28.1* | PPA20393 | RS4139 | *tu1859* | 19bp insertion | exon 3 | This paper |
| *Ppa-daf-22.1;Ppa-daf-22.2* | ppa_stranded_DN16812_c0_g1_i1;PPA41516 | RS2770 | *tu489;tu504* | 7bp deletion;7bp insertion | refer to Markov et al., 2016 | Markov et al., 2016 |
| *sult-1* | PPA12547 | RS2974 | *tu1061* | 10bp deletion | exon 9 | Namdeo et al., 2018 |
| *nag-1;nag-2* | PPA06134;PPA34489 | RS3195 | *tu1142;tu1143* | 1 SNP + 17bp insertion;2 SNPs + 9bp deletion | exon 5;exon 5 | Sieriebriennikov et al., 2018 |
